# Ephrin-B2 Paces Neuronal Production in the Developing Neocortex

**DOI:** 10.1101/2020.01.10.901587

**Authors:** Anthony Kischel, Christophe Audouard, Mohamad-Ali Fawal, Alice Davy

## Abstract

**Background:** During mammalian cerebral cortex development, different types of projection neurons are produced in a precise temporal order and in stereotypical numbers. The mechanisms regulating timely generation of neocortex projection neurons and ensuring production in sufficient numbers of each neuronal identity is only partially understood.

**Results:** Here, we show that ephrin-B2, a member of the Eph:ephrin cell-to-cell communication pathway, sets the neurogenic tempo in the neocortex. Indeed, conditional mutant embryos for ephrin-B2 exhibit a transient delay in neurogenesis and acute stimulation of Eph signaling by in utero injection of synthetic ephrin-B2 led to a transient increase in neuronal production. Using genetic approaches we show that ephrin-B2 acts on neural progenitors to control their differentiation in a juxtacrine manner. Unexpectedly, we observed that perinatal neuron numbers recovered following both loss or gain of ephrin-B2, highlighting the ability of neural progenitors to adapt their behavior to the state of the system in order to produce stereotypical numbers of neurons.

**Conclusions:** Altogether, our data uncover a role for ephrin-B2 in embryonic neurogenesis and emphasizes the plasticity of neuronal production in the neocortex.

## BACKGROUND

It is estimated that the mouse brain is composed of about 80 million neurons, the vast majority of which are generated during fetal life. These neurons come in dozens of different flavors, each neuronal subtype being characterized by wiring partners, location, morphology, neurotransmitter production. The mechanisms ensuring that neuron types and numbers are faithfully produced are only partly understood. Over the years this question has been largely studied in the neocortex, a mammalian-specific region of the brain which is the siege of complex cognitive functions.

Neurogenesis in the neocortex is characterized by the orderly production of different types of projection neurons that migrate and settle at precise spatial positions, thus eventually forming a six-layered structure. These neurons can be grouped in two main classes: early born deep layer (DL) neurons which project to subcortical targets and late born upper layer (UL) neurons that extend inter-hemispheric projections [1]. Both DL and UL neurons are born from neural progenitors (NP) located in the germinal zones of the neocortex, namely the ventricular zone (VZ) and the subventricular zone (SVZ). At the origin of these NP are neuroepithelial cells that actively proliferate to amplify the initial pool of NP and then transition to the apical progenitor (AP) fate. At the onset of neurogenesis there are two main types of NP in the neocortex: Pax6+ AP whose soma are located in the VZ and Tbr2+ basal progenitors (BP) populating the SVZ. Lineage tracing studies in the mouse have shown that asymmetric division of AP give rise mainly to BP and to ~10% of neurons while BP give rise to 90% of cortical neurons [2–4]. More recently, a new type of BP, outer radial glial cells (oRG), has been identified and shown to be massively expanded in the primate neocortex compared to rodents [5]. Because these cells are able to undergo a large number of symmetric divisions before terminal differentiation, they are believed to be the cellular source for the massive expansion of the neocortex in primates [6].

Time is a key parameter in neocortex development, influencing both the number and the type of neuron generated [7]. Indeed, the duration of the initial NP expansion phase, the rate of NP self-renewal vs. differentiation, as well as the dynamics of NP division are important temporal parameters that have impacts on final neuron numbers. One striking illustration of this is the difference in pace of development that distinguishes mouse and human neocortex development. Neuron production lasts about a week in the mouse vs. 17 weeks in humans and this correlates with an estimated thousand fold increase in neuronal outputs [8]. In addition, timing of differentiation defines the type of neuron produced, with sequential production of DL neurons first and UL neurons last. While mechanisms controlling NP self-renewal vs differentiation have been studied in great detail, mechanisms that are crucial to a timely and orderly neuronal production are less characterized.

Accumulating evidence indicates that progression in the temporal sequence relies on information provided to NP by their local environment. The first observations suggesting that extrinsic information was important to set the pace of neurogenesis in the neocortex came from heterochronic transplantation studies in the ferret showing that environmental cues present at late stages of cortical development within the germinal zone can “re-program” a young NP to give rise to a late-born projection neurons [9]. This question has been reinvestigated with modern techniques recently and it was shown that late AP remain temporally plastic and can adapt to earlier temporal environments by changing their molecular and neurogenic fate [10]. In contrast, late born BP lack this temporal plasticity [10]. Several studies in the mouse have identified extrinsic signals, mainly in the form of feedback from post mitotic neurons, that act on NP to modulate the orderly production of projection neurons [11–15]. More strikingly, it has been recently demonstrated that neuronal loss induced from early to mid corticogenesis is compensated by the over production of UL neurons at late stages of corticogenesis in the mouse [16], indicating that NP are able to sense and respond to environmental cues, adapting their behavior to modulate neuronal production either in types or in numbers.

Eph/ephrin signaling is a bi-directional cell-to-cell communication pathway that plays important roles in tissue patterning [17] but has also been implicated in neurogenesis mainly in the adult brain [18]. In vitro and in vivo studies in the adult mouse have shown that one member of the family, ephrin-B2, is a neurogenic cue [19, 20], however the role of this membrane-bound protein in embryonic neurogenesis in the neocortex has not been reported. Here, we hypothesized that ephrin-B2 could play a role in the timely production of projection neurons in the neocortex. Using gain and loss of function paradigms in vivo we describe a role for ephrin-B2 in pacing neuronal production in the neocortex. At mid-corticogenesis, neuronal numbers are decreased in loss of function conditions and increased in gain of function conditions. Unexpectedly, we show that both in loss and gain of function situations, neuronal numbers are normal at birth.

## RESULTS

### Ephrin-B2 is expressed in progenitors and modulates neuronal production

To address a potential role for ephrin-B2 in controlling neurogenesis in the developing neocortex, we first surveyed its expression in this tissue by in situ hybridization (ISH), from the neuroepithelial stage (E10.5) to the peri-natal stage (E18.5) (Figure 1A). *Efnb2* is strongly expressed in neuroepithelial cells at E10.5 and it remains expressed in NP at E13.5. At E13.5, expression of *Efnb2* is also detected in the cortical plate (CP), in a high-lateral to low-medial gradient which coincides with the progression of neurogenesis. At later stages, expression of *Efnb2* is low in progenitors and in DL neurons, while high expression is observed in UL neurons. To assess expression of *Efnb2* in NP at single cell resolution, we made use of a reporter mouse line that expresses a nuclear GFP under the control of the endogenous *Efnb2* promoter [21]. Epifluorescence detection of GFP in thick vibratome sections of the neocortex at E12.5 shows that *Efnb2* is expressed in the majority of NP and is strongly upregulated in new born neurons located basally to the VZ (Figure 1B). Co-immunostaining with an antibody that detects the phosphorylated form of EphB1-3 indicates that these receptors are phosphorylated both in NP and in neurons (Figure 1B) suggesting that EphB:ephrinB2 signaling is active in these cells. To uncover the functional significance of this activation, we generated conditional mutant embryos using *Efnb2*^*floxed*^ [22] mice and the *Nestin-Cre* allele [23] which fully excises *Efnb2* as early as E11.5 in the neocortex as shown by in situ hybridization (Sup Figure 1A). First, to evaluate the consequence of deleting *Efnb2* on Eph:ephrin signaling we monitored the phosphorylation status of EphB1-2 in the neocortex of E13.5 control and *Efnb2*^*lox/lox*^; *Nestin-Cre* (cKO^Nes^) embryos. Western blot analysis shows that tyrosine phosphorylation of EphB1-3 is decreased in the conditional mutants (Figure 1C). In parallel, we monitored the phosphorylation status of EphA4, which is also a cognate receptor for ephrin-B2. No change in the phosphorylation status of EphA4 was observed in cKO^Nes^ embryos (Figure 1C). Altogether, these results indicate that loss of ephrinB2 specifically impairs EphB signaling in the developing neocortex.

**Figure 1.**
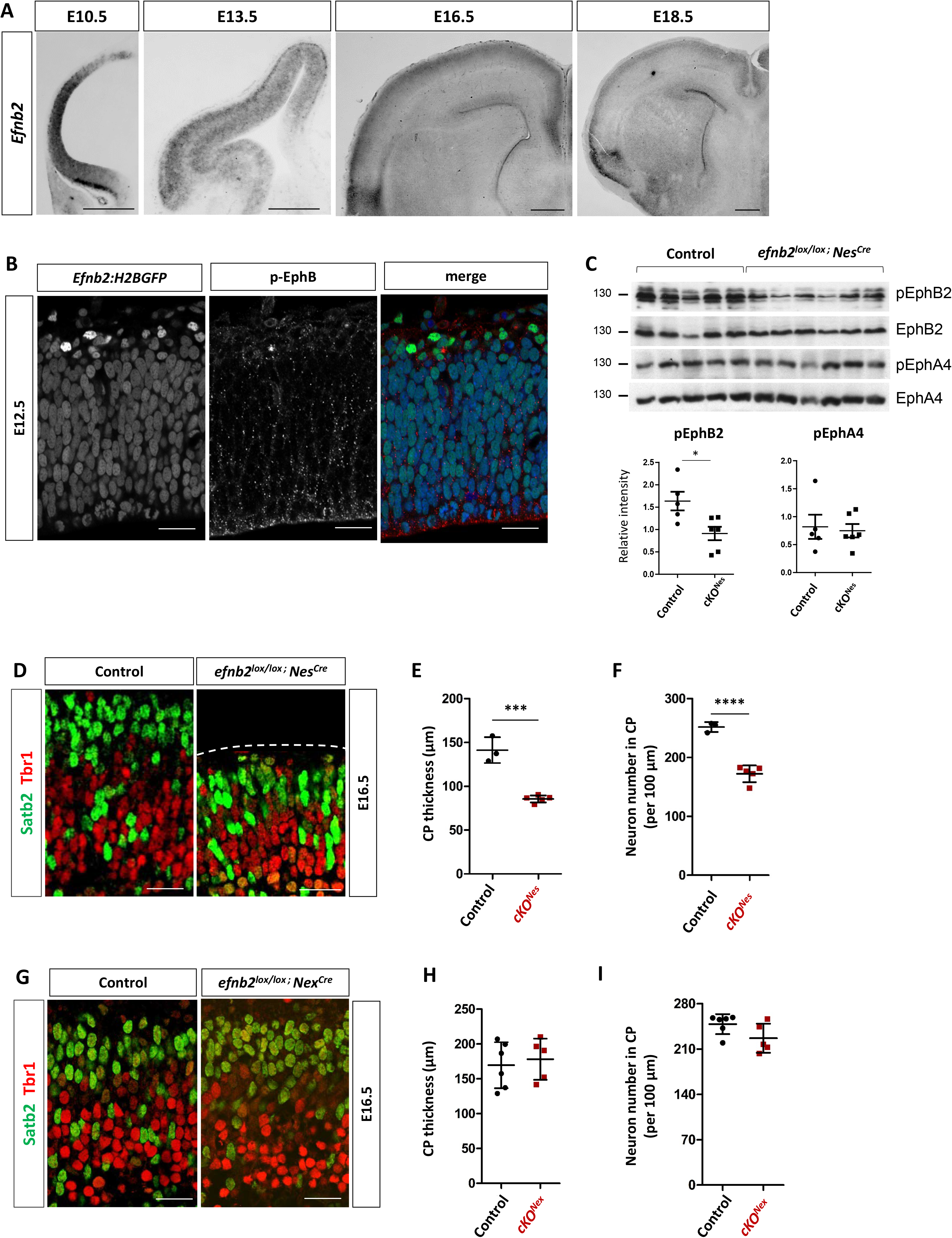
Ephrin-B2 is dynamically expressed in the developing neocortex and modulates neuronal production. A. *Efnb2* in situ hybridization on transverse sections of the neocortex at different developmental stages (indicated). Scale bar: 500 μm. B. Epifluorescence (GFP; green) detection on a transverse section of the neocortex of an E12.5 *Efnb2:H2BGFP* embryo. The section was immunostained with a phospho-EphB antibody (red). C. Western blot analysis of E13.5 neocortex tissue extracted from control (n= 5) and *Efnb2*^*lox/lox*^; *Nes-Cre* (n= 6) embryos. Antibodies are indicated on the side. Quantification of the signal (bottom) shows the ratio between pEph and total Eph. D. Transverse sections of the cortical plate of the neocortex of E16.5 control and *Efnb2*^*lox/lox*^; *Nes-Cre* embryos were immunostained for Tbr1 (red) and Satb2 (green). E. Measurement of cortical plate (CP) thickness in control (n= 3) and *Efnb2*^*lox/lox*^; *Nes-Cre* (n= 5) embryos. The data is the average of lateral, central, medial measurements. F. Quantification of the number of neurons in the cortical plate of the neocortex. The graph represents total neuron numbers (Tbr1+ and Satb2+ neurons) in control (n= 3) and *Efnb2*^*lox/lox*^; *Nes-Cre* (n=5) embryos. The data is the average of lateral, central, medial counts. G. Transverse sections of the cortical plate of the neocortex of E16.5 control and *Efnb2*^*lox/lox*^; *Nex-Cre* embryos were immunostained for Tbr1 (red) and Satb2 (green). H. Measurement of cortical plate thickness in control (n= 5) and *Efnb2*^*lox/lox*^; *Nex-Cre* (n= 5) embryos. The data is the average of lateral, central, medial measurements. I. Quantification of the number of neurons in the cortical plate of the neocortex. The graph represents total neuron numbers (Tbr1+ and Satb2+ neurons) in control (n= 5) and *Efnb2*^*lox/lox*^; *Nex-Cre* (n=5) embryos. The data is the average of lateral, central, medial counts. Data are reported as mean ± SEM (***P < 0.001; ****P < 0.0001). CP: cortical plate. Scale bars B-G: 50 μm.

Next we analyzed neuronal numbers at E16.5 in control and cKO^Nes^ embryos using a combination of Tbr1 and Satb2 antibodies that are markers for early born and late born neurons, respectively. This combination of antibodies labels the majority of neocortical projection neurons during neocortex development. Immunostaining and imaging of paraffin sections revealed an apparent decrease in the number of neurons in the CP of cKO^Nes^ embryos compared to controls (Figure 1D). To quantify this phenotype, we measured the CP thickness (Figure 1 E) and manually counted neuron numbers (Figure 1F) in control and cKO^Nes^ embryos. Both parameters were reduced in cKO^Nes^ embryos indicating that excision of *Efnb2* leads to a reduction in neuron numbers in the neocortex CP. Of note, we observed that the reduction in neuron numbers was mostly due to a decrease in Satb2+ neurons and that it followed a mediolateral gradient, with a stronger reduction medially than laterally (Sup Figure 2A, B). Importantly, the decreased number of neurons in the CP of cKO^Nes^ embryos did not correlate with Satb2+ cells stacked in the intermediate zone, in fact the intermediate zone surface area was reduced (Sup Figure 2C, D) nor with an increased number of apoptotic cells (Sup Figure 2E) ruling out cell death or migration defects as potential causes for the observed phenotype.

As described above, *Efnb2* is expressed both in neurons and in progenitors and the *Nestin-Cre* allele eliminates *Efnb2* expression in both populations. To ask whether neuronal or progenitor ephrin-B2 expression is required to control neuron numbers, we generated conditional mutant embryos using the *Nex-Cre* mouse line [24] which excises *Efnb2* in neurons but not in progenitors (cKO^Nex^) as shown by ISH (Sup Figure 1B). We performed immunostaining with Tbr1 and Satb2 on paraffin sections of E16.5 embryos (Figure 1G) and measured the thickness of the CP and quantified neuronal numbers. No difference in CP thickness (Figure 1H) nor in the number of neurons (Figure 1I) was observed in the cKO^Nex^ mutant embryos. Altogether, these results indicate that ephrin-B2 is required in neocortical progenitors to control the production of Tbr1+ and Satb2+ neurons.

### The decrease in neuron numbers in cKO^Nes^ embryos is transient

The fact that the reduction in neuron numbers followed a mediolateral gradient at E16.5 hinted at a dynamic evolution of the phenotype. Hence, to better characterize the phenotype observed in cKO^Nes^ embryos we quantified CP thickness and neuron numbers throughout corticogenesis by imaging region of interest (ROI) on tissue sections. We performed immunostaining with Tbr1 and Satb2 at two early stages of corticogenesis and 2 perinatal stages (Figure 2A). These analyses showed that while CP thickness is significantly decreased at E16.5 in cKO^Nes^ embryos, by E18.5 the decrease in no longer statistically significant and at P4 there is no detectable difference (Figure 2B). With respect to neuron numbers, no statistically significant difference was observed in cKO^Nes^ at E13.5 but at E14.5 and E16.5 the number of neurons was decreased in cKO^Nes^ (Figure 2C). This decrease is no longer observed at E18.5 and P4 (Figure 2C). To capture the evolution of neuronal numbers in a dynamic way, and to ask whether early born and late born neurons were equally or differentially affected, we plotted the mean neuronal numbers for each type of neurons at each developmental stages and represented the data as curves (Figure 2D-F and Sup Figure 3). It is important to note that the method of 2D quantification has limitations when comparing different developmental stages since it does not take into consideration the 3D expansion of the tissue, nor the decrease in cell density at late stages due to neuronal arborization. Thus, this approach only provides an estimate for neuronal number dynamics. However, this approach is completely appropriate to compare genotypes at each stage. Altogether, these results clearly show that loss of ephrin-B2 leads to a decrease in neuronal production at early to mid-stages of corticogenesis that is compensated at late stages of neocortex development. Interestingly, early born Tbr1+ neurons tend to be mostly decreased at E14.5 (without reaching statistical significance) and their number is compensated at E16.5 (Figure 2E and Sup Figure 3B, C) while late born Satb2+ neurons are severely diminished at E16.5 and their number are compensated at E18.5 (Figure 2F and Sup Figure 3C, D). Altogether, this data indicates that loss of ephrin-B2 transiently impairs the production of both types of neurons, but with a stronger effect on Satb2+ neurons.

**Figure 2.**
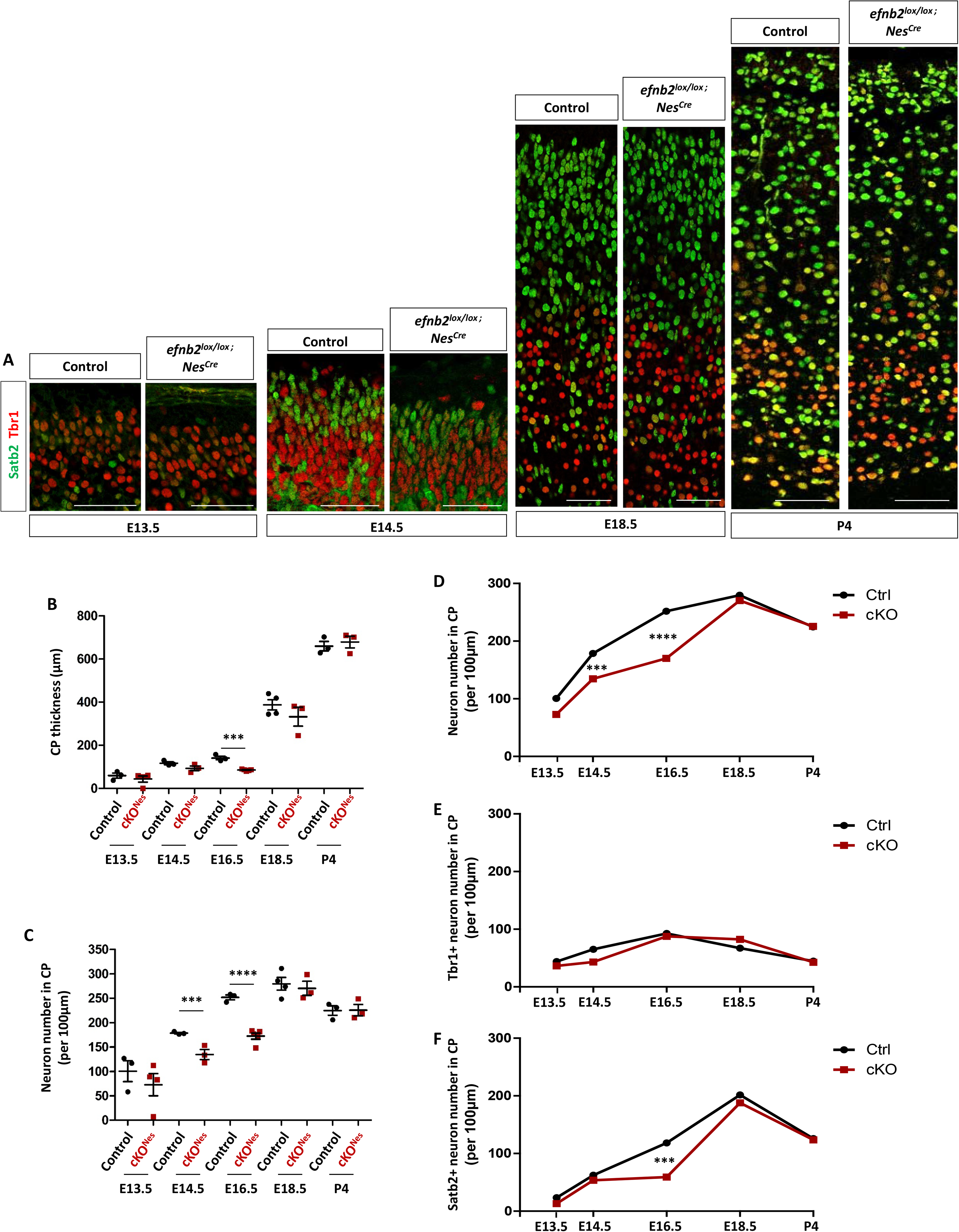
The decrease in neuronal number in the neocortex of *Efnb2* cKO^Nes^ is transient. A. Transverse sections of the cortical plate of the neocortex of control and *Efnb2*^*lox/lox*^; *Nes-Cre* embryos at different developmental stages (indicated) were immunostained for Tbr1 (red) and Satb2 (green). B. Measurement of cortical plate thickness in control (n=3 or 4) and *Efnb2*^*lox/lox*^; *Nes-Cre* (n=3 or 4) embryos at all stages. C. Quantification of the number of neurons in the cortical plate of the neocortex at all stages. The graph represents total neuron numbers (Tbr1+ and Satb2+ neurons) in control and *Efnb2*^*lox/lox*^; *Nes-Cre* embryos. D. Mean total neuron numbers at each developmental stages of both genotypes were plotted on a curve graph. E. Mean numbers of Tbr1+ neurons at each developmental stage were plotted on a curve graph. F. Mean numbers of Satb2+ neurons at each developmental stage were plotted on a curve graph. Data are reported as mean ± SEM (***P < 0.001; ****P < 0.0001). CP: cortical plate. Scale bars: 50 μm.

### Decreased neuron numbers in cKO^Nes^ embryos correlate with slight changes in progenitor numbers

Decreased neuronal production could be caused by various processes affecting progenitors such as changes in cell cycle duration, modifications in progenitor numbers, imbalance in self-renewal vs. differentiation. To discriminate between these possible causes, we performed double EdU/BrdU labelling as previously described [25] to estimate progenitor cell cycle length and to quantify progenitor populations. To capture progenitor cell cycle length we combined EdU/BrdU immunostaining with Tbr2 immunostaining and considered that Tbr2+ cells are BP and Tbr2-cells in the VZ are AP (Figure 3A). We manually counted the number of progenitors (Tbr2-VZ cells and Tbr2+ cells called P cells), the number of EdU+ P cells (called L cells) and the number of EdU+BrdU+ P cells (called S cells) in control and cKO^Nes^ E13.5 embryos (see also Methods and Sup Figure 4A, B). These numbers were used to estimate cell cycle length as described previously [25]. No difference in cell cycle length was observed between control and cKO^Nes^ embryos when analyzing AP and BP progenitors together (Figure 3B) or separately (Sup Figure 4C). Next, we re-analyzed the data to quantify numbers of each type of progenitors in both genotypes. With this analysis we detected a slight but statistically significant increase in the total number of progenitors in cKO^Nes^ embryos which does not appear to be restricted to one type of progenitors (Figure 3C). A similar trend was observed at E14.5 without reaching statistical significance (Figure 3D, E). These data suggest that modification in cell cycle length is unlikely to account for the neuronal phenotype observed in cKO^Nes^ embryos. Instead, these results suggest that loss of ephrin-B2 could impair the differentiation process and favor the progenitor fate.

**Figure 3.**
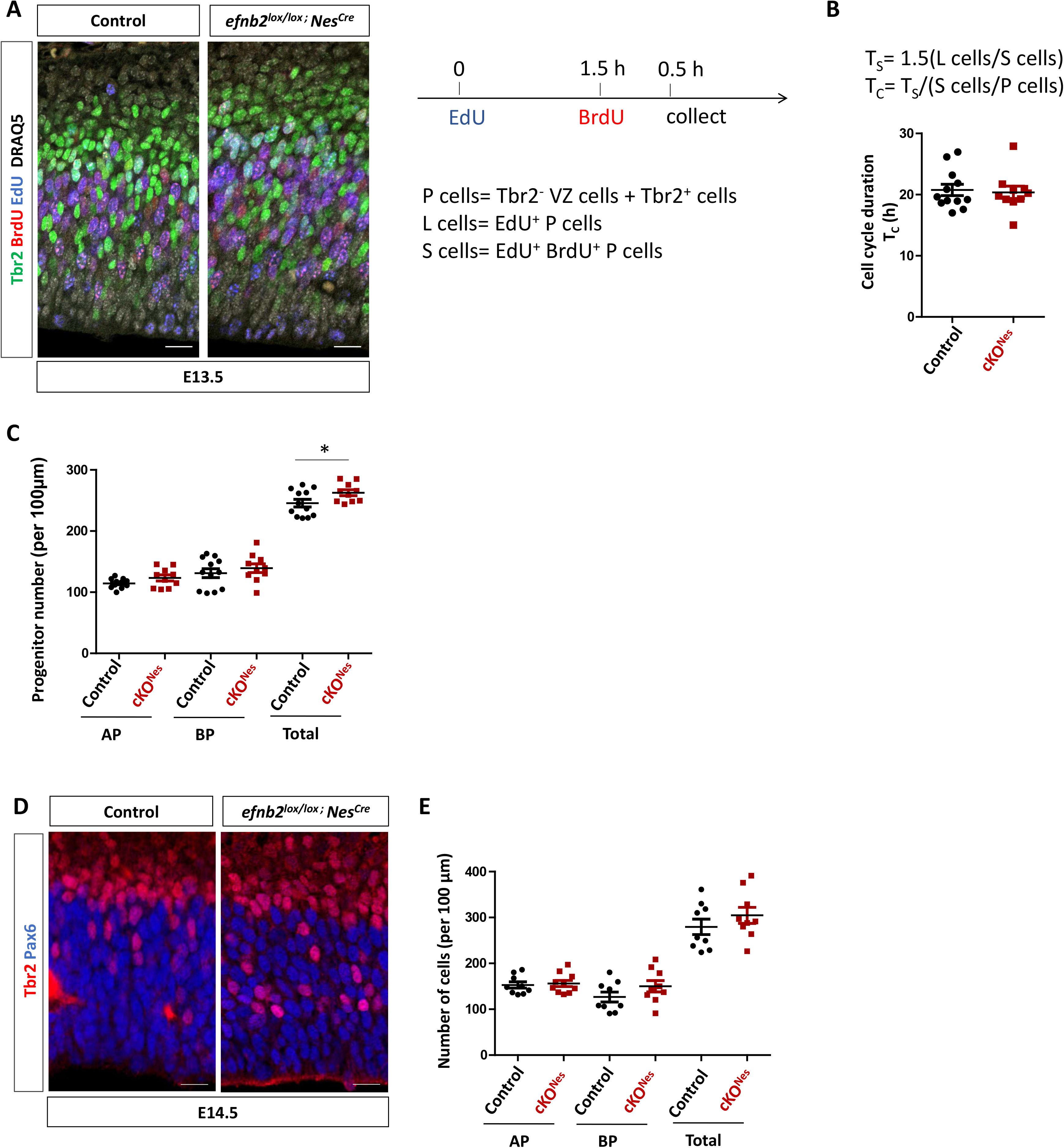
Cell cycle duration and numbers of progenitors in *Efnb2* cKO^Nes^. A. Transverse sections of the neocortex of EdU/BrdU-injected E13.5 control and *Efnb2*^*lox/lox*^; *Nes-Cre* embryos were immunostained for BrdU (red), Tbr2 (green) and EdU (blue). Scale bars: 25 μm. The injection protocol and the various cell populations that were counted are indicated on the right. B. S-phase duration (T_s_) and cell cycle duration (T_c_) were calculated using the indicated formulas. The graph shows T_c_ in control (n=6) and *Efnb2*^*lox/lox*^; *Nes-Cre* embryos (n=5). Data for each counted hemisphere are plotted. C. Quantification of AP (Tbr2-), BP (Tbr2+) and total progenitor numbers in control (n=6) and *Efnb2*^*lox/lox*^; *Nes-Cre* embryos (n=5). Data for each counted hemisphere and mean ± SEM are reported, unpaired t-test with Welch’s correction (*P < 0.05). D. Transverse sections of the neocortex of E14.5 control and *Efnb2*^*lox/lox*^; *Nes-Cre* embryos were immunostained for Tbr2 (red) and Pax6 (blue). Scale bars: 25 μm. E. Quantification of AP (Pax6+), BP (Tbr2+) and total progenitor numbers in control (n=4) and *Efnb2*^*lox/lox*^; *Nes-Cre* embryos (n=4). Mean ± SEM are reported.

### Injection of eB2-Fc in lateral ventricles leads to a transient increase in neuronal production

To test whether ephrin-B2 directly acts as a signal to promote neuronal differentiation (and thus neuronal production), we injected clustered recombinant ephrin-B2-Fc protein (eB2-Fc) or EphB2-Fc protein into the lateral ventricles of developing E13.5 embryos to artificially (and transiently) activate Eph forward or ephrin reverse signaling, respectively. Injected brains were collected at E15.5 and E18.5 and tissue sections were immunostained with Tbr1 and Satb2 antibodies (Figure 4A). Quantification of neuron numbers showed that a single injection of eB2-Fc (but not EphB2-Fc; Sup Figure 5) leads to an increased number of neurons at E15.5 compared to control injected samples (Figure 4B). Interestingly, at E18.5 control and eB2-Fc injected samples had similar numbers of neurons (Figure 4B) indicating that the tissue is able to adapt to both too few or too many neurons. Detailed examination of the data by neuron populations reveal that Satb2+ neurons account for most of the increase in neuronal production at E15.5 (Figure 4C), again suggesting that the production of this population of neurons is highly sensitive to the ephrin-B2 signal. The increased neuronal production at E15.5 correlated with a decreased number of Pax6+ AP in eB2-Fc injected samples (Figure 4D). These results indicate that transient activation of Eph forward but not ephrin reverse signaling in AP promotes neuronal production and they reveal that the tissue is able to compensate both mid-corticogenesis decrease and increase in neuronal numbers.

**Figure 4.**
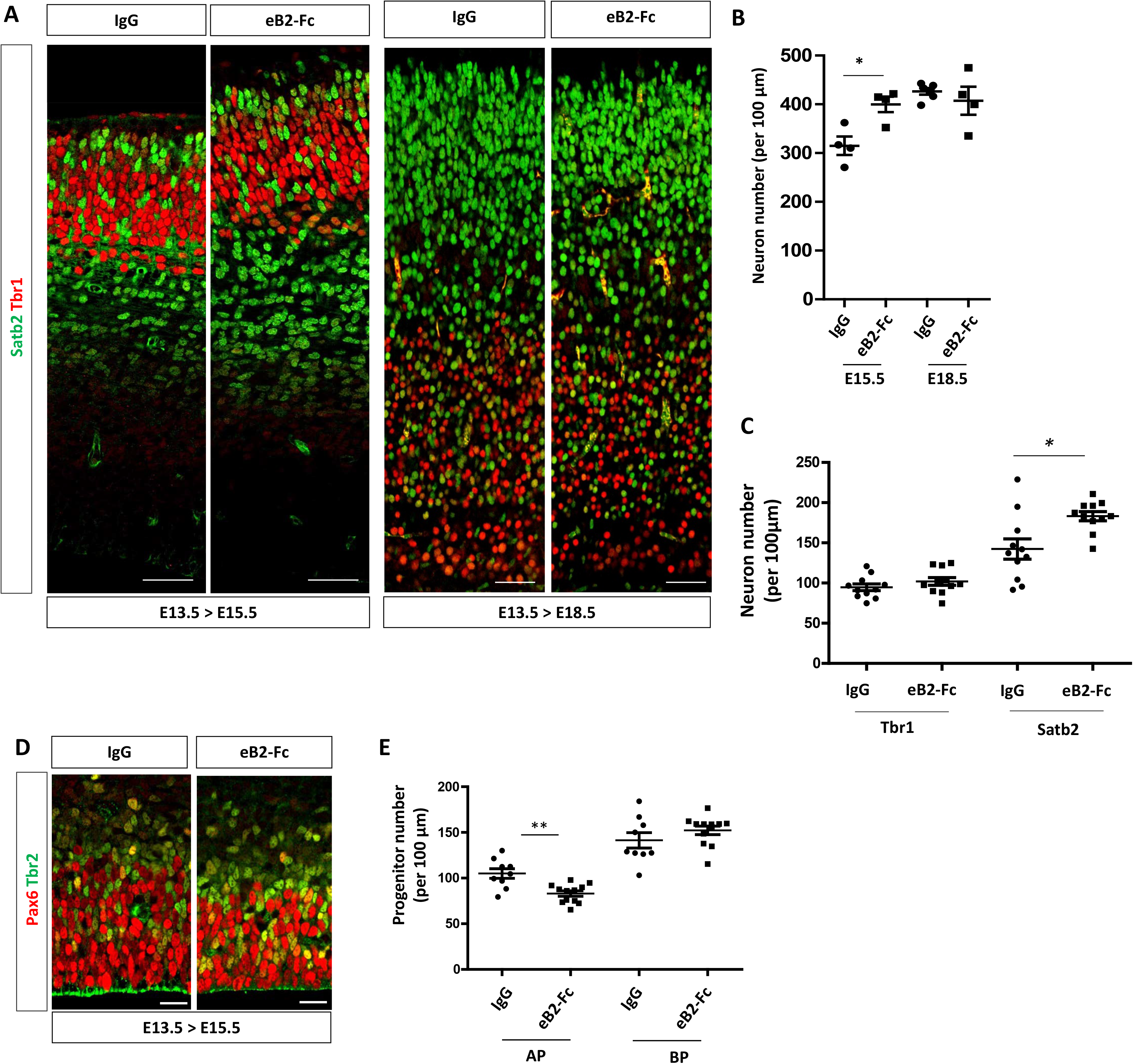
Acute injection of eB2-Fc in the lateral ventricle leads to an increased neuronal production. A. Transverse sections of the neocortex of E15.5 or E18.5 control injected embryos (IgG) (n=4) and embryos injected with eB2-Fc (n=4) were immunostained for Tbr1 (red) and Satb2 (green). Scale bars: 50 μm. B. Quantification of the number of neurons in the neocortex of E15.5 and E18.5 embryos. The graph represents total neuron numbers (Tbr1+ and Satb2+ neurons in the lateral region of the neocortex) in control and eB2-Fc injected embryos. C. Quantification of the number of Tbr1+ and Satb2+ neurons in the neocortex of E15.5 injected embryos (central region of the neocortex). Data for each counted hemisphere are plotted. D. Transverse sections of the ventricular zone and subventricular zone of E15.5 control injected embryos (IgG) and embryos injected with eB2-Fc (central region of the neocortex) were immunostained for Pax6 (red) and Tbr2 (green). Scale bars: 20 μm. E. Quantification of the number of Pax6+ and Tbr2+ progenitors in IgG (n=4) and eB2-Fc (n=5) injected embryos. Data for each counted hemisphere are plotted. Data are reported as mean ± SEM. Mann-Whitney statistical test (*P < 0.05; **P < 0.01).

## DISCUSSION

### Ephrin-B2 is a pro-neurogenic cue in the embryonic neocortex

Our study reveals an unappreciated role for ephrin-B2 in controlling neuronal production in the embryonic neocortex. We show that conditional excision of ephrin-B2 leads to a transient reduction in neuronal numbers while a single injection of clustered eB2-Fc recombinant protein in lateral ventricles of developing embryos induces a transient increase in neuronal production. This is reminiscent to the role of ephrin-B2 in promoting neurogenesis in the adult hippocampus which has been reported previously [20]. Indeed, hippocampal bilateral injection of clustered eB2-Fc recombinant proteins led to an increased commitment of NSC to the neuronal fate at the expense of the neural stem cell fate while loss of ephrin-B2 decreased neurogenesis in the subgranular zone. One interesting aspect of that study was to show that ephrin-B2 was expressed and required in astrocytes to control neural stem cell commitment in a juxtacrine manner [20]. Here, we show that ephrin-B2 is expressed and required in NP but our data does not discriminate between a requirement in AP vs. BP. However, the injection experiments suggest that ephrin-B2 acts on NP in a juxtacrine manner to activate Eph signaling and to promote neurogenesis. Although the gain and loss of function data are consistent in terms of neuronal production, the observed changes at the level of NP are different. In the loss of function experiment, both populations of Tbr2- and Tbr2+ NP appear to be slightly affected while in the gain of function experiment it is the population of Pax6+ AP that is modified. This discrepancy could be due to the fact that AP are primarily targeted in the injection paradigm while in the genetic loss of function paradigm, ephrin-B2 is excised from both AP and BP. Of note, we showed recently that a decrease in Sox2+ NP and an increase in neuron numbers is also observed following injections of clustered eB1-Fc in lateral ventricles of the developing mouse neocortex [26]. Whether ephrin B1 and ephrin-B2 promote neuronal production via similar mechanisms will have to be explored further.

The onset of neuronal production does not appear to be affected in absence of ephrin-B2, suggesting that the impairment in neuronal production in *Efnb2* cKO^Nes^ at mid-corticogenesis is due to a slowing down of the neurogenic process rather than a delay. What could be the cause of this? Based on a similar phenotype observed in *Brap* mutant embryos, we hypothesized that cell cycle duration might be changed. Indeed, loss of Brap, a Ras-Erk signaling modulator with E3 ligase activity led to a prolonged cell cycle which impeded neuronal production without significantly affecting the number of Pax6+ and Tbr2+ NP [27]. However, using the sequential EdU/BrdU injection protocol we were not able to detect a difference in cell cycle duration in E13.5 cKO^Nes^ embryos, indicating that cell cycle alteration is unlikely to be the cause of the neuronal phenotype. Instead, our data suggest that ephrin-B2 controls the transition from AP to BP to neurons. Indeed, in both loss and gain of function conditions we observed an inverse correlation between progenitor numbers and neuron numbers. It is puzzling, however that the observed changes in progenitor numbers were very small compared to changes in neuron numbers.

### A compensatory mechanism normalizes neuron numbers in the neocortex

The fact that neuronal production is compensated over time in the neocortex of *Efnb2* mutants highlights the existence of a compensatory mechanism during corticogenesis. It also suggests that ephrin-B2 is required transiently, at early stages of corticogenesis, a time window that matches the peak of its expression in NP. A similar phenotype has been shown for Caspr (Contactin-associated protein), a member of the Neurexin family of adhesion molecules. Similar to ephrin-B2, Caspr expression peaks at early stages of corticogenesis in the ventricular zone and Caspr mutant exhibit transient decrease in neuronal production which was compensated at E18.5 [28]. The author proposed that Caspr controlled the timing of NP differentiation by modulating Notch signaling [28]. A similar powerful compensation mechanism was also evidenced recently by transiently inducing neuronal cell death in the neocortex using genetic tools. In this genetic context, the loss of DL neurons was compensated by an over production of UL neurons, which, the authors showed, resulted from the increased proliferation of BP [16]. Altogether, these studies highlight the fact that compensation is possible when early and transient regulatory mechanisms are impacted. On the other hand, many genes that cause autosomal recessive primary microcephaly encode centrosomal proteins that are ubiquitously expressed [29] and this may explain why no compensation is observed when these genes are mutated. Simlary, Brap mutants exhibit microcephaly at P28 [27], indicating that neuronal production is not compensated in these animals, probably because Brap is a ubiquitously-expressed protein whose function is required in NP throughout corticogenesis.

In the future, it would be interesting to identify external signals and intrinsic pathways that allow NP to sense the state of neuronal production and adapt their behavior (e.g. self-renewal vs proliferation vs differentiation). Clearly, our data shows that these compensatory mechanisms are not dependent on ephrin-B2. One candidate could be the neurotrophin Ntf3 which is produced by early born neurons, and acts on NP to promote the switch from AP to BP [12]. It is important to note that while this role of Ntf3 was convincingly described using in utero electroporation experiments, *Ntf3*^*-/-*^ mutant neocortex exhibit only a mild neuronal phenotype at E18.5 [12], again suggesting compensatory mechanisms. Another mechanism could be direct Notch signaling from newborn neurons to progenitors via retention of their apical endfoot [15]. Mechanical cues could also represent potential sensing mechanisms. Indeed, tissue crowding or mechanical stretch have been shown to regulate cell proliferation in other contexts [30] and mechanical morphogenesis is now being recognized as a driver of neocortex development and evolution [31]. Another important question that will have to be addressed in the future is whether neurons that are generated in an inappropriate time-window upon compensation are fully functional and correctly integrated into neuronal circuits. This is particularly relevant in the context of human studies which have shown that *EfnB2* haploinsufficicency causes neurodevelopmental delay [32].

## CONCLUSIONS

Our study adds ephrin-B2 to the list of extrinsic signals that control neurogenesis and it adds embryonic neuronal production to the list of ephrin-regulated functions. Further, it highlights that development of the neocortex is plastic, since this tissue is able to adapt its neuronal output to the state of the system.

## METHODS

### Animals

Mice were bred in the animal facility. *Efnb2*^*H2BGFP*^, *Efnb2*^*loxlox*^, *Nestin-Cre* and *Nex-Cre* mouse lines have been described previously [21–24]. To generate conditional mutants, *Efnb2*^*loxlox*^ females were bred with *Nestin-Cre; Efnb2*^*loxlox*^ males. Neonates were individually genotyped by PCR.

### Primary antibodies

Satb2: Abcam, Mouse, ab51502, Paraffin: 1/150, Vibratome: 1/40.

Tbr1: Abcam, Rabbit, ab31940, Paraffin: 1/150, Vibratome: 1/100.

Pax6: Covance, Rabbit, PRB-278P, 1/300.

Tbr2: Invitrogen, Rat, 14-4875-82, 1/100.

Cleaved Caspase 3: Cell signaling, Rabbit, 9661, 1/300.

Eph B2: R & D Systems, Goat, AF467, WB: 1/1000.

Eph A4: Cell signaling, Rabbit, 8793, WB: 1/2000.

P-Eph B1-2: Abcam, Rabbit, ab61791, WB: 1/1000.

P-Eph A4: Gift from Greenberg’s lab, Harvard medical school, Rabbit, WB: 1/1000.

### In situ hybridization

In situ hybridization was performed on transverse vibratome sections. Briefly, embryos were fixed 24 hours in 4% paraformaldehyde (PFA), sectioned on the vibratome and sections were dehydrated in ethanol. Following rehydration, sections were treated with proteinase K (10 μg/ml in PBS/0.1% Tween-20) for 7 minutes at room temperature and subsequently post-fixed in PFA/glutaraldehyde solution. Sections were incubated overnight at 65°C in hybridization buffer (50% formamide, 5x SSC (pH 6), 0.1% SDS, 50 μg/ml heparin, 500 μg/ml yeast RNA) containing the labelled probe. Sections were washed twice with solution I (5x SSC, 50% formamide, 0.1% SDS) at 65°C and 3 times in solution III (2x SSC, 50% formamide, 0.1% SDS) at 65°C, rinsed in TBS/0.1% Tween-20 and incubated overnight in blocking buffer (TBS with 2% goat serum, 0.1% blocking reagent (Roche), 0.1% Tween-20) containing an AP-labelled anti-DIG antibody (1/2000) (Roche). NBT/BCIP was used as a substrate for the Alkaline Phosphatase. For double labelling, in situ hybridization was followed by post-fixation (4% PFA for 30 minutes) and immunostaining was performed as described above.

### Protein Extraction and Western Blot

For protein extraction, neocortex was dissected from E13.5 embryos in ice-cold PBS (Cat#D1408, Sigma) solution. Whole cell extracts were obtained by re-suspending tissues in the following ice-cold lysis buffer (150 mM NaCl (Cat#1112-A, Euromedex), 50 mM Tris-HCl pH 7.4 (Cat#EU0011, Euromedex), 0.5 mM EDTA (Cat#EU0007, Euromedex), 2 mM Na3VO4(Cat#S6508, Sigma), 1% Nonidet P-40 (NP-40) (Cat#N6507, Sigma), 0.5 mM EGTA (Cat#E4378, Sigma) and 0.1 mM PMSF (Cat#78830, Sigma)) supplemented by protease inhibitors (Cat#**11836170001, Roche)** for 1h on ice. Protein lysates were then vortexed, sonicated with Biorupter® Sonicator for 10 cycles (Cat#B01020001, Diagenode), and centrifuged at 13,000 rpm for 10 min. Cleared lysates were either used for Western blot analyses. Samples were denatured by boiling in loading buffer (4× 100 mM Tris-HCL, pH 6.8, 8% SDS (Cat#EU0660, Euromedex), 40% glycerol (Cat#G9012, Sigma), 4% β-mercaptoethanol (Cat#63689, Sigma), and bromophenol blue (Cat#B0126, Sigma)) before loading and electrophoresis on an 8 or 16% SDS-PAGE gel. Proteins were transferred onto a nitrocellulose membrane (Cat#**GE10600002**, GE Healthcare), which was blocked for 30 min and incubated with primary antibody in 5% nonfat dry milk in TBS-T (20 mM Tris base (Cat#26-128-3094-B, Euromedex), 150 mM NaCl, and 0.05% Tween 20 (Cat#2001-A, Euromedex) adjusted to pH 7.6 with 1 M HCl (Cat#30721, Sigma)) overnight at 4°C. Milk was replaced with 5% BSA (Cat#04100811C, Euromedex) for detection of phosphorylated epitopes.

### In utero injections

Timed-pregnant mice were anesthetized using Vetflurane (Cat#Vnr137317, Vibrac) and uterine horns were exposed. 1µg/ml human IgG or 1µg/ml pre-clustered eB2-Fc (Cat#476-EB, R&S Systems) were injected in the lateral ventricle of E13.5 embryos neocortex using pulled micropipettes. Body wall cavity and skin were sutured and embryos were allowed to develop normally for 48h or until E18.5. Embryonic brains were collected at E15.5 or E18.5, fixed with 4% paraformaldehyde (Cat#15710, Fisher scientific) overnight at 4°C. Vibratome sections (50 µm) (Cat#VT1000 S, Leica Biosystems) were processed for immunofluorescence staining as described below.

### Immunofluorescence

Timed-pregnant mice were sacrificed by cervical dislocation; embryos were removed and dissected in ice-cold PBS. For immunohistochemistry, embryonic brain tissues were collected and placed in 4% PFA at 4ºC. All brain samples were removed from PFA and either equilibrated in 70% ethanol and embedded in paraffin or immediately sectioned on a vibratome. Coronal paraffin sections (5 μm) were obtained and placed on Superfrost microscope slides (Fisher Scientific) and stored at room temperature until use.

Vibratome sections were permeabilized and blocked with PBTA2 solution (2% BSA, 2% FBS, 1% Tween20 (Cat#2001-A, Euromedex) for 2h at room temperature. Primary antibodies diluted in PBTA2 and secondary antibodies diluted in PBS were incubated about 16h at 4°C and 1.5h at room temperature respectively. Labelled sections were mounted on glass slides (Superfrost) with Coverslips using mounting medium (4.8% wt/vol Mowiol and 12% wt/vol glycerol in 50 mM Tris, pH 8.5).

Paraffin sections were rehydrated and demasking was performed by incubating slides in pH 6 Citrate buffer for 45min at 90°C. Immunostaining was then performed as described above.

### Image acquisition and quantification

Microscope acquisitions were performed on an inverted SP8 (40x, TCS; Leica Biosystems) and entire hemispheres were acquired for each embryo. Observation was performed using Type-F mineral oil (1153859; Leica Biosystems). Cell counting was performed with the ImageJ software. To standardize cell counting, for each developmental stage, sections at similar rostral positions were selected, using anatomical landmarks (shape and size of the LGE, shape of the lateral ventricle, presence of the choroid plexus). For the loss of function experiments, at developmental stages E13.5 and E14.5, a single region of interest (lateral) per hemisphere was used for manual counting. At later stages, 3 regions of interest (lateral, central and medial) were used for manual counting. In this case, the data presented in the graphs is the average of the 6 regions of interest (2 hemispheres) for each embryo. For the gain of function experiments, a single region of interest (lateral or central) per hemisphere was used for counting.

### Cell cycle analyses

We used the protocol described by Martynoga [25]. Briefly, pregnant dam (E13.5) were injected with a single dose (100 μl) of EdU (10 μg/ml), followed 1h30 later by a single dose (100 μl) of BrdU (10 μg/ml). Embryos were collected 30 minutes later and processed for tissue sectioning and immunostaining to detect EdU+ cells, BrdU+ cells and Tbr2+ cells. We manually counted Tbr2-cells in the VZ corresponding to Pax6+ AP and Tbr2+ cells in a region of interest encompassing the central region of the neocortex. It is important to note that the EdU/BrdU double labeling method can be used to estimate cell cycle length only when analyzing populations of cycling cells, yet, Tbr2 remains expressed in newborn neurons which are no longer cycling. Thus, to circumvent this caveat we estimated the fraction of Tbr2+ cells that also express Tbr1 in both genotypes analyzed (Sup Figure 4) and assumed that these cells represent newborn neurons that have exited the cell cycle. We then used this to adjust the total number of Tbr2+ progenitors and obtain the number of P cells. We next counted L cells (P cells that are EdU+) and S cells (P cells that are EdU+BrdU+). Length of the S-phase and length of the cell cycle were calculated using these numbers, as described in Figure 3B and [25].

### Statistical analyses

For experiments involving a single pair of conditions, statistical significance between the two sets of data were analyzed with an unpaired-t-test (Mann-Whitney) with Prism5 (GraphPad software). For datasets containing more than two samples, one-way analysis of variance (ANOVA) was used. Sample sizes of sufficient power were chosen on the basis of similar published research. For neuronal numbers, each dot on a graph represents a different embryo for which, depending on the stage, 3 ROI were manually counted, on 3-4 different sections. For progenitor numbers, each dot on a graph represent a single ROI averaged from 3 different sections for a single hemisphere. Between 3-6 embryos for each experimental conditions were analyzed. The value on the graph is the mean of these 3-4 points. Statistically significant differences are reported at *P < 0.05, **P < 0.01, ***P < 0.005, ****P < 0.001.

## DECLARATIONS

### Ethics approval

Animal procedures were approved by the appropriate Animal Care Committee (APAFIS#1289-2015110609133558 v5).

### Authors’ contributions

AK and AD conceptualized the study, AK designed, performed and analyzed experiments and AD wrote the manuscript. MAF performed western-blot experiments. CA performed in utero injections and immunofluorescence experiments on paraffin sections. All authors contributed to analyzing experiments and to the editing of the manuscript.

## Acknowledgements

We thank Alain Vincent, Eric Agius and members of the Davy and Pituello teams for helpful discussions and comments on the manuscript. We acknowledge core support from the Imagery Platform of Toulouse and from the Anexplo animal facility.

