## Supplemental Figures for "Ephrin-B2 Paces Neuronal Production in the Developing Neocortex"

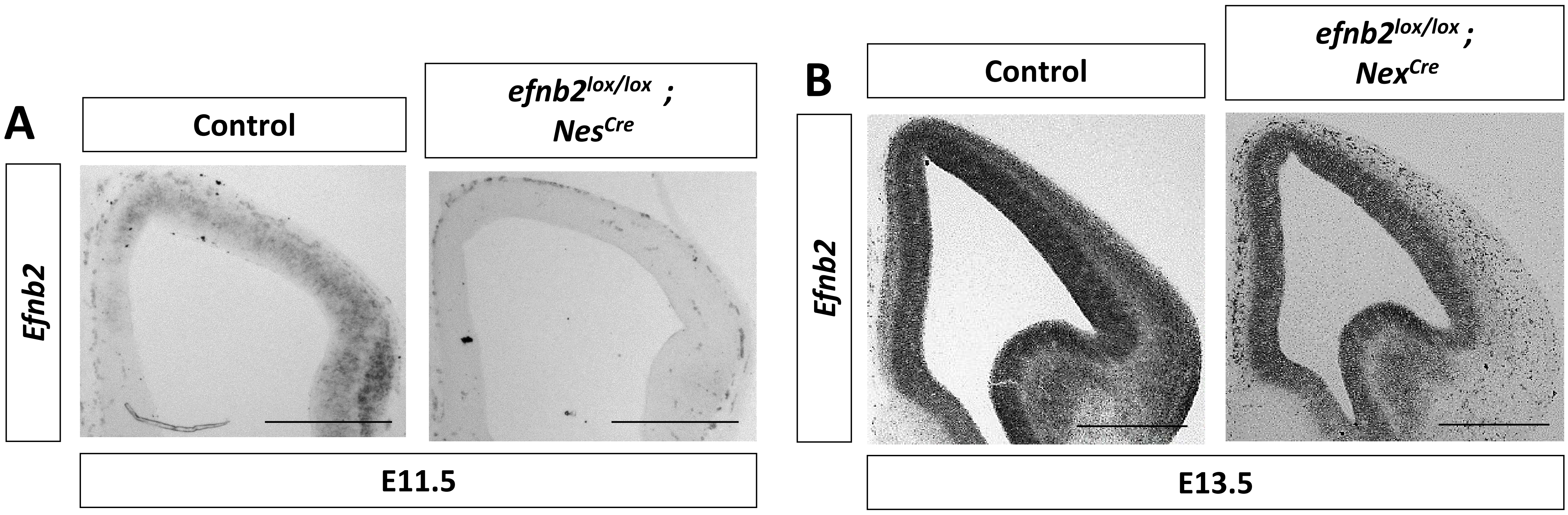


**Sup Figure 1. Validation of lox-Cre excision of *Efnb2.***

A. *Efnb2* in situ hybridization on transverse sections of the neocortex of E11.5 control and *Efnb2*$^{lox/lox}$*; Nes-Cre* embryos. Scale bar: 500 mm.

B. *Efnb2* in situ hybridization on transverse sections of the neocortex of E13.5 control and *Efnb2*$^{lox/lox}$*; Nex-Cre* embryos. Scale bar: 500 mm.

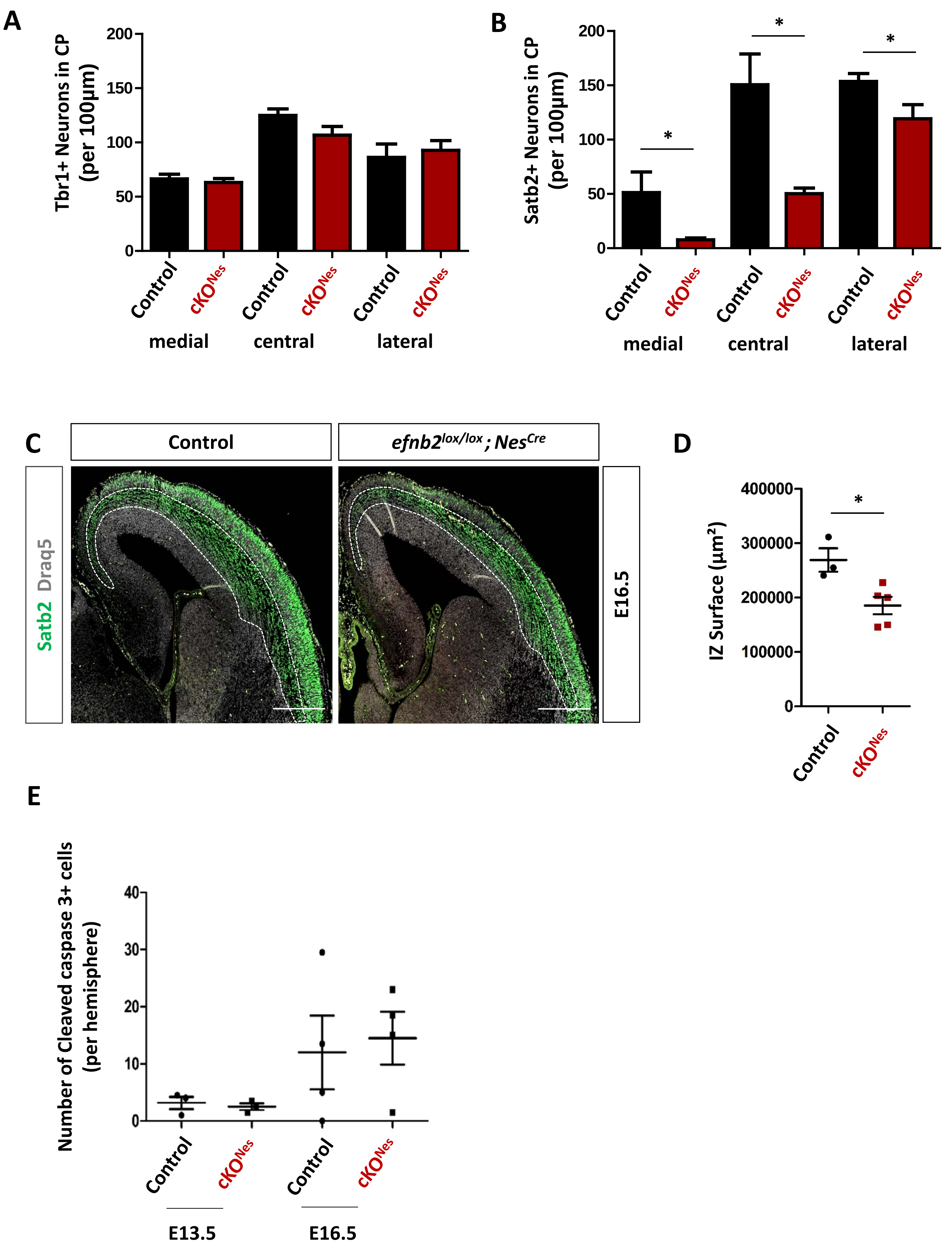


**Sup Figure 2. Decreased numbers of neurons in *Efnb2* mutants is not due to defective migration or increased apoptosis.**

A. Quantification of the number of Tbr1+ neurons in medial, central and lateral regions of the cortical plate of the neocortex in control (n= 3) and *Efnb2^lox/lox ; Nes-Cre* (n=5) E16.5 embryos.

B. Quantification of the number of Satb2+ neurons in medial, central and lateral regions of the cortical plate of the neocortex in control (n= 3) and *Efnb2^lox/lox ; Nes-Cre* (n=5) E16.5 embryos.

C. Transverse sections of the neocortex of E16.5 control and *Efnb2^lox/lox ; Nes-Cre* embryos were immunostained for Satb2 (green) and stained with Draq5 (grey). The intermediate zone was delimited based on DAPI staining (white outline).

D. Quantification of the IZ surface area of control (n=3) and *Efnb2^lox/lox ; Nes-Cre* (n=5) embryos.

E. Quantification of the number of cleaved caspase3+ cells per hemisphere in control and *Efnb2^lox/lox ; Nes-Cre* embryos at two different developmental stages (indicated).

Data are reported as mean ± SEM (*P < 0.05). IZ : intermediate zone. Scale bar represents 500 mm.

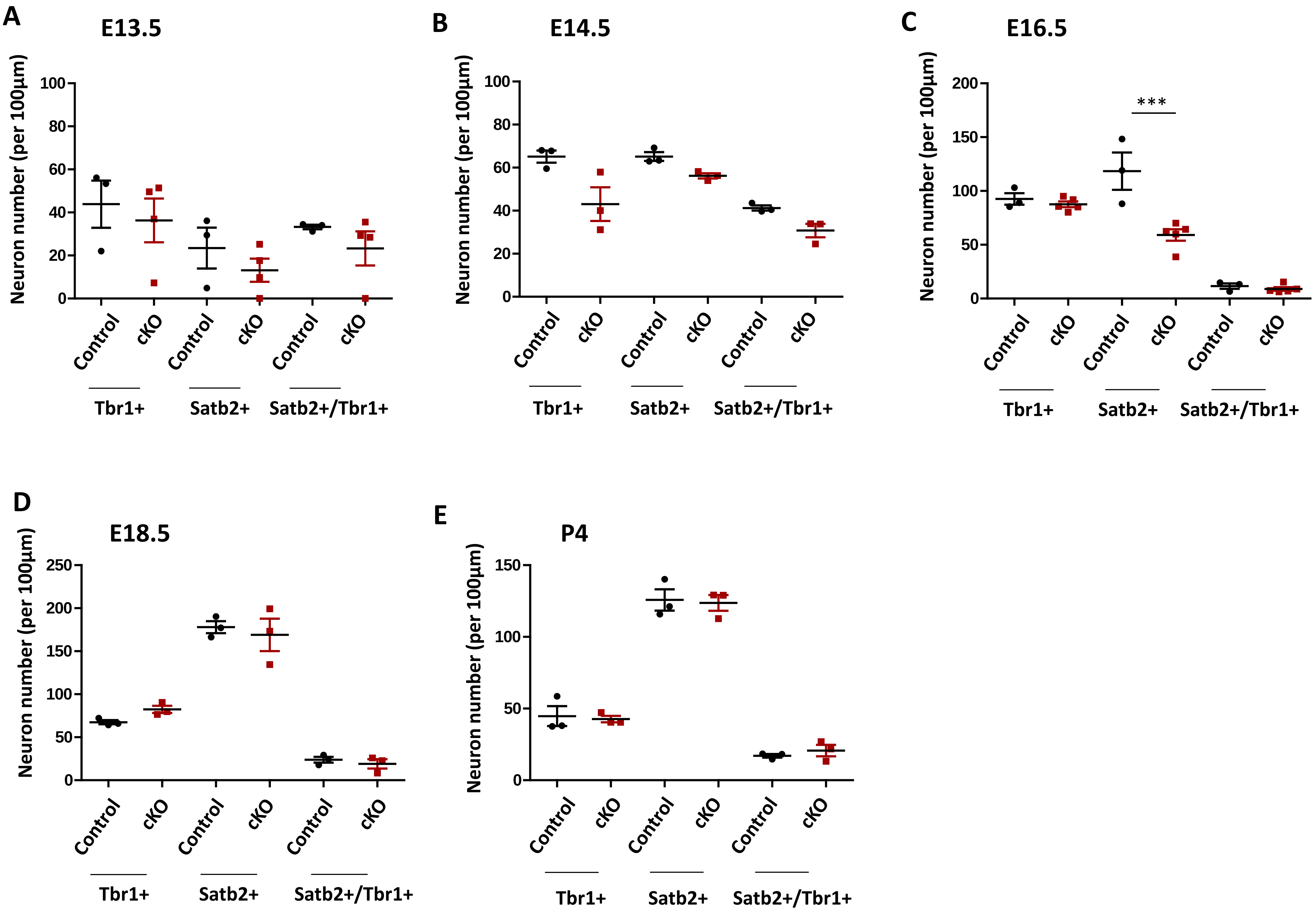


**Sup Figure 3. Quantification of neuron numbers by type (Tbr1+ and Satb2+) at different developmental stages.**

A-E. Quantification of Tbr1+ and Satb2+ neuron number at each developmental stage in both genotypes. At E13.5, the data is highly variable from embryo to embryo. At E14.5, the statistically significant decrease in total neuron numbers is due to a non statistically significant decrease in both Tbr1+ and Satb2+ neurons while at E16.5, only Satb2+ neurons are decreased, suggesting that the number of Tbr1+ neurons has been compensated by this stage.
Data are reported as mean ± SEM, Mann-Whitney statistical test (***P < 0.001).

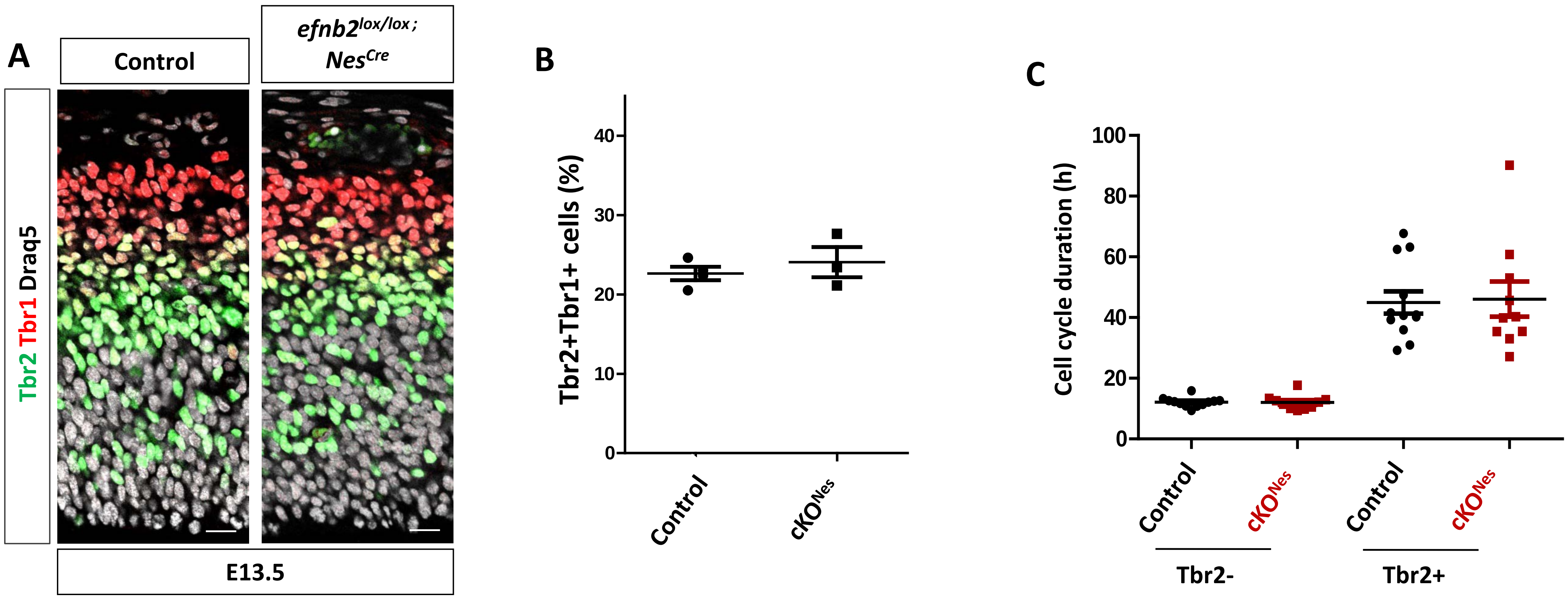


**Sup Figure 4. Cell cycle analyses.**

A. Transverse sections of the neocortex of E13.5 control and *$Efnb2^{lox/lox}$ ; Nes-Cre* embryos were immunostained for Tbr2 (green), Tbr1 (red) and stained with Draq5 (grey).
B. Quantification of the fraction of Tbr2+ cells that express Tbr1 in E13.5 control and *$Efnb2^{lox/lox}$ ; Nes-Cre* embryos (n=3).
C. Estimation of cell cycle length of Tbr2- and Tbr2+ progenitors in control (n=6) and *$Efnb2^{lox/lox}$ ; Nes-Cre* (n=5) E13.5 embryos. The data for Tbr2+ progenitors is highly variable in both genotypes underlying the difficulty of using this method for this population of progenitors.

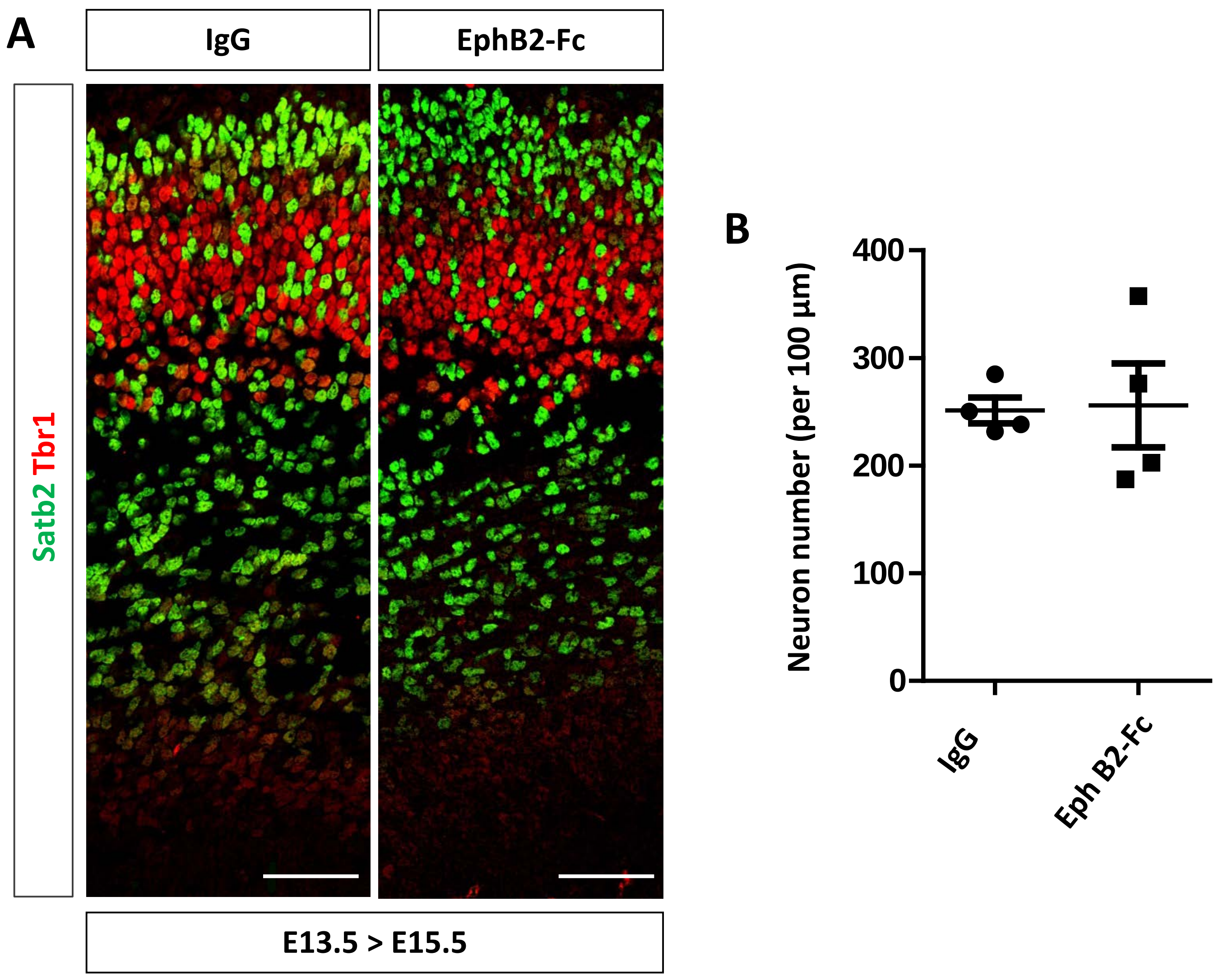


**Sup Figure 5. Injection of EphB2-Fc in the lateral ventricle does not modify neuron numbers.**

A. Transverse sections of the neocortex of E15.5 control injected embryos (IgG) (n=4) and embryos injected with EphB2-Fc (n=4) were immunostained for Tbr1 (red) and Satb2 (green). Scale bars: 50 mm.

B. Quantification of the number of neurons in the neocortex of E15.5 embryos. The graph represents total neuron numbers (Tbr1+ and Satb2+ neurons in lateral, central and medial regions of the neocortex) in control and EphB2-Fc injected embryos. Data are reported as mean ± SEM.
